# Transcriptome-wide identification of coding and noncoding RNA-binding proteins defines the comprehensive RNA interactome of *Leishmania mexicana*

**DOI:** 10.1101/2021.12.08.471363

**Authors:** Karunakaran Kalesh, Wenbin Wei, Brian S. Mantilla, Theodoros I. Roumeliotis, Jyoti Choudhary, Paul W. Denny

**Affiliations:** Department of Chemistry, Durham University, Durham, United Kingdom; Department of Biosciences, Durham University, Durham, United Kingdom; Functional Proteomics Group, The Institute of Cancer Research, London, United Kingdom

**Keywords:** RNA-binding proteins, *Leishmania*, OOPS, TMT labelling, LC-MS/MS, RNA-seq, non-Poly(A) interactome, WDR domain

## Abstract

Proteomic profiling of RNA-binding proteins in *Leishmania* is currently limited to polyadenylated mRNA-binding proteins, leaving proteins that interact with nonadenylated RNAs, including noncoding RNAs and pre-mRNAs, unidentified. Using a combination of unbiased orthogonal organic phase separation methodology and tandem mass tag-labelling-based high resolution quantitative proteomic mass spectrometry, we robustly identified 2,417 RNA-binding proteins, including 1289 putative novel non-poly(A)-RNA-binding proteins across the two main *Leishmania* life cycle stages. Eight out of twenty *Leishmania* deubiquitinases including the recently characterised *L. mexicana* DUB2 with an elaborate RNA-binding protein interactome were exclusively identified in the non-poly(A)-RNA-interactome. Additionally, an increased representation of WD40 repeat domains were observed in the *Leishmania* non-poly(A)-RNA-interactome, thus uncovering potential involvement of this protein domain in RNA-protein interactions in *Leishmania*. We also characterise the protein-bound RNAs using RNA-sequencing and show that in addition to protein coding transcripts ncRNAs are also enriched in the protein-RNA interactome. Differential gene expression analysis revealed enrichment of 145 out of 195 total *L. mexicana* protein kinase genes in the protein-RNA-interactome, suggesting important role of protein-RNA interactions in the regulation of the *Leishmania* protein kinome. Additionally, we characterise the quantitative changes in RNA-protein interactions in hundreds of *Leishmania* proteins following inhibition of heat shock protein 90 (Hsp90). Our results show that the Hsp90 inhibition in *Leishmania* causes widespread disruption of RNA-protein interactions in ribosomal proteins, proteasomal proteins and translation factors in both life cycle stages, suggesting downstream effect of the inhibition on protein synthesis and degradation pathways in *Leishmania*. This study defines the comprehensive RNA interactome of *Leishmania* and provides in-depth insight into the widespread involvement of RNA-protein interactions in *Leishmania* biology.

**IMPORTANCE:** Advances in proteomics and mass spectrometry have revealed the mRNA-binding proteins in many eukaryotic organisms, including the protozoan parasites *Leishmania* spp., the causative agents of leishmaniasis, a major infectious disease in over 90 tropical and sub-tropical countries. However, in addition to mRNAs, which constitute only 2 to 5% of the total transcripts, many types of non-coding RNAs participate in crucial biological processes. In *Leishmania*, RNA-binding proteins serve as primary gene regulators. Therefore, transcriptome-wide identification of RNA-binding proteins is necessary for deciphering the distinctive posttranscriptional mechanisms of gene regulation in *Leishmania*. Using a combination of highly efficient orthogonal organic phase separation method and tandem mass tag-labelling-based quantitative proteomic mass spectrometry, we provide unprecedented comprehensive molecular definition of the total RNA interactome across the two main *Leishmania* life cycle stages. In addition, we characterise for the first time the quantitative changes in RNA-protein interactions in *Leishmania* following inhibition of heat shock protein 90, shedding light into hitherto unknown large-scale downstream molecular effect of the protein inhibition in the parasite. This work provides insight into the importance of total RNA-protein interactions in *Leishmania*, thus significantly expanding our knowledge of the emergence of RNA-protein interactions in *Leishmania* biology.

---

Unicellular parasites of the genus *Leishmania* are the causative agents of leishmaniasis, a major vector-borne infectious disease in 98 tropical and sub-tropical countries (1). Being evolutionarily ancient eukaryotes *Leishmania* spp. possess several distinctive biological features, such as lack of transcription factors, polycistronic transcription and RNA trans-splicing (2). Consequently, the gene expression in *Leishmania* is exclusively regulated by posttranscriptional mechanisms such as RNA processing, degradation, and protein translation (2). RNA-binding proteins (RBPs) in *Leishmania* serve as *trans*-regulators and control the processing and trafficking of RNA molecules from synthesis to degradation (3). Therefore, identification of the complete repertoire of the RBPs is considered a prerequisite for deciphering the mechanisms of gene regulation in *Leishmania* spp. A technique known as RNA Interactome Capture (RIC) that relies on a selective binding interaction of poly(A) tails in mature mRNAs with oligo(dT) beads is widely used for the extraction of mRNA-binding proteins in various organisms including *Leishmania* spp. (3-13). Despite its success, RIC is limited to proteins that bind only mature mRNAs bearing poly(A) tails. However, non-poly(A) RNA molecules including noncoding RNAs (ncRNAs) comprise the major portion of the total transcribed molecules in the cell. In addition to the mRNAs the ncRNAs also function as ribonucleoprotein particles (RNPs) and carry out biological functions including synthesis of new proteins, RNA processing, genome remodelling and regulation of transcription (14, 15). We therefore envisaged a comprehensive transcriptome-wide identification of coding and non-coding RBPs in the *Leishmania* spp. Towards this we applied the recently reported orthogonal organic phase separation (OOPS) method (16, 17) in combination with tandem mass tag (TMT) labelling-based quantitative proteomic mass spectrometry (MS) (18) and report herein the most comprehensive identification of RBPs in *Leishmania mexicana* (*L. mexicana*) parasites to date.

Many advancements in the identification of RBPs have recently occurred. By showing that the more RBPs a protein interacts with the more likely that protein itself is an RBP, a computational approach termed SONAR (support vector machine obtained from neighbourhood associated RBPs), could predict RBPs in any organism (19). Although SONAR does not rely on structural homology or prior knowledge of RNA-binding domains, availability of large-scale protein-protein interaction (PPI) data set is often limited in many organisms. Also, available PPI data sets may not have covered all RBPs. Recent experimental developments in the capturing of RBPs include enhanced-RIC (e-RIC), a modification of the RIC protocol that uses a locked nucleic acid modified capture probe to mitigate the nonspecific binding problem of the original RIC method (20), metabolic incorporation of nucleotide analogues (21, 22), and the use of organic-aqueous phase separation-based methods namely the OOPS (16), protein cross-linked RNA extraction (XRNAX) (23) and phenol-toluol extraction (PTex) (24). Although powerful the e-RIC is still limited to poly(A)-binding RBPs. The phase separation-based methods use UV cross linking (CL) to generate protein-RNA adducts which are isolated from free protein and free RNA by at least one round of acidic phenol phase partitioning. The OOPS employs multiple rounds of phase partitioning to enrich the RBPs at an acidic guanidinium thiocyanate-phenol-chloroform (AGPC) interface. In this case, RNase treatment of the enriched interface followed by a final AGPC phase partitioning enables release of the RBPs into the organic layer.

Whilst both the nucleotide analogue-based methods and the phase separation-based methods enable capturing of coding and noncoding RBPs the former is largely limited to RBPs that bind nascent RNAs as prolonged treatment of the nucleotide analogues will inhibit rRNA synthesis, and compromise cell viability (25, 26). We therefore employed the OOPS method for the capturing of total RBPs in the *Leishmania* parasite. The OOPS protocol for the discovery of total RBPs in mammalian cell lines employed stable isotope labeling by amino acids in cell culture (SILAC) (27) to compare the relative abundance of proteins in the cross-linked (CL) and non-cross-linked (NC) samples (16). In the case of *Leishmania*, however, culturing of the parasites in the SILAC media resulted in growth defects in our hands (unpublished results). We therefore employed the highly sensitive peptide-level TMT-labelling and high-resolution quantitative proteomic MS to robustly compare the CL and NC OOPS samples in both promastigote and axenic amastigote *L. mexicana* life cycle stages. This study significantly expands the RBP landscape of *Leishmania*. Furthermore, we showed that the classical heat shock protein 90 (Hsp90) inhibitor tanespimycin regulates the RNA-binding property of hundreds of *L. mexicana* RBPs, shedding light into hitherto unknown large-scale downstream molecular effect of this small molecule inhibitor in the parasite.

## RESULTS

### Assessment of OOPS protocol in *L. mexicana* promastigotes and axenic amastigotes

First, we performed the OOPS protocol at varying UV-doses in both *L. mexicana* promastigotes and axenic amastigotes (Fig. 1A). We quantified the RNA recovered from the interface following protein digestion using proteinase K and extraction from the aqueous phase of a final AGPC phase partition. As in the case of mammalian cells reported earlier (16), we observed UV-dose-dependent migration of the RNA from the aqueous phase to the interface, saturating at around 500 mJ/cm^2^ and at approximately 75% and 70% respectively of the total RNA content in promastigote and axenic amastigote life cycle stages of the parasite (Fig. 1B and C; Supplementary Fig. S1). This gave us the confidence to couple the OOPS with large-scale quantitative proteomic MS to characterise the total RBPs in the *L. mexicana* parasites.

**FIG. 1.**
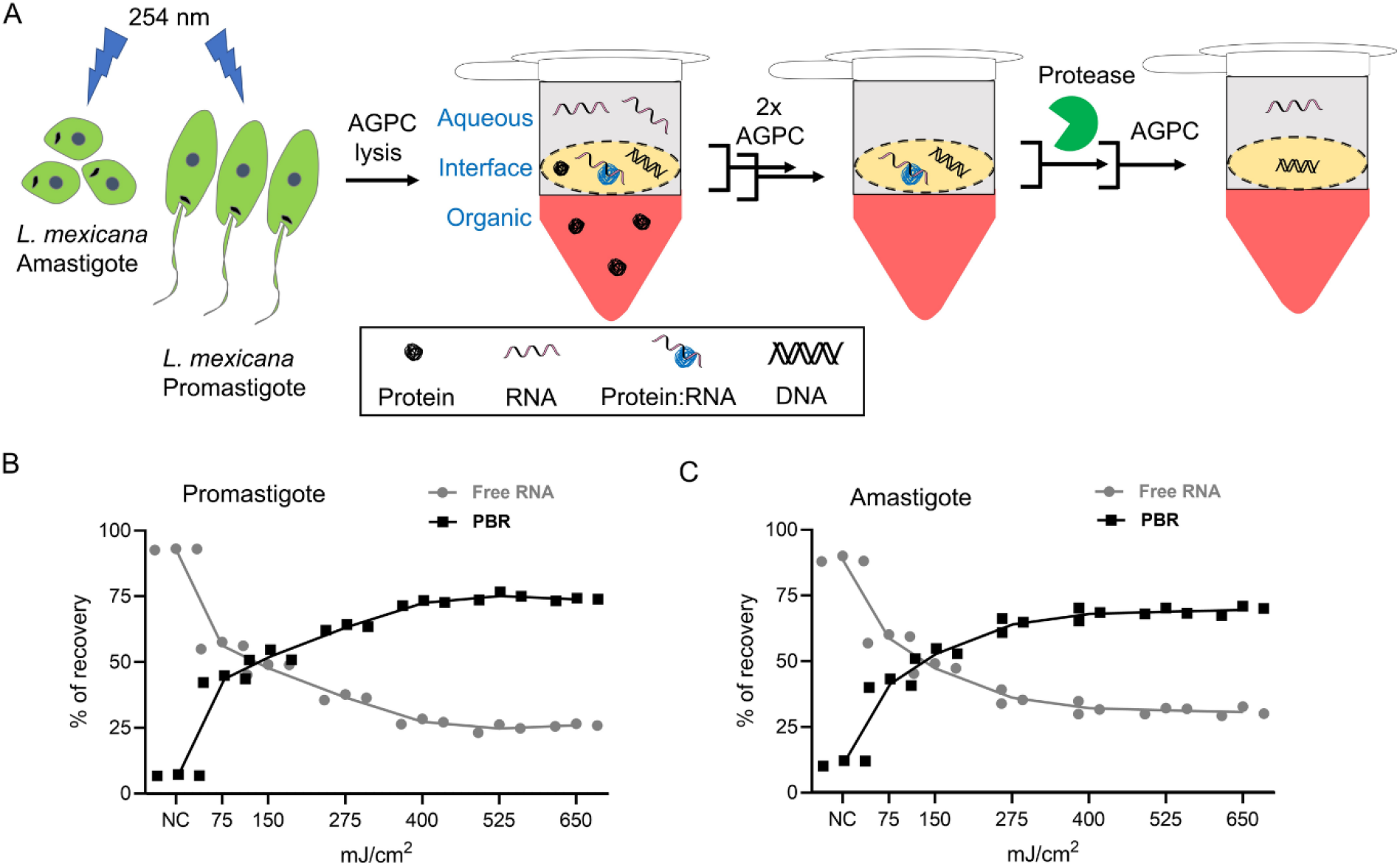
OOPS recovers protein-bound RNAs (PBRs) in *Leishmania*. (A) Schematic representation of the OOPS method to extract PBRs in *L. mexicana* promastigotes and axenic amastigotes. UV-irradiation of the live *Leishmania* parasites induce RNA-protein cross-linking. Upon cell lysis and AGPC phase-partitioning, the protein-RNA adducts, which have physical properties of both proteins and RNAs, are simultaneously drawn to the organic and aqueous phases and therefore accumulate in the interface. Sequential AGPC phase-partitioning followed by protease digestion of the interface and a final AGPC separation yields the previously PBRs in the aqueous phase. Relative proportions of free-RNAs (aqueous phase) and PBRs (interface) with increasing UV-dosage in promastigotes (B) and in axenic amastigotes (C). Data shown as mean ± s.d. of three independent experiments. NC: non-cross-linked controls.

### Capture of total RBPs in *L. mexicana* promastigotes and axenic amastigotes using OOPS

Next, in order to comprehensively identify the coding and non-coding RBPs in both promastigote and amastigote life cycle stages of the *Leishmania* parasites, we developed a workflow by combining the OOPS with tandem mass tag (TMT) labelling-based quantitative proteomic MS (Fig. 2A). We performed OOPS with a fixed UV-dose of 525 mJ/cm^2^ and without UV-irradiation in three independent experiments for promastigotes and axenic amastigotes and the RBPs in the interfaces were recovered into the organic phases by a final AGPC phase partition following RNase treatment. Proteins, after digestion with trypsin and TMT-sixplex labelling, were analysed by nano-LC-MS/MS. With a filter minimum of 2 unique peptides and a requirement of minimum 6 valid values across the three replicates of the UV-irradiated and non-irradiated conditions for the promastigotes and axenic amastigotes, a total of 2417 RBPs were identified in the OOPS samples (Table S1a). This includes 1255 proteins in the promastigotes (Table S1b) and 1960 proteins in the amastigotes (Table S1c) identified with 2 or more unique peptides. The OOPS-TMT-sixplex combination revealed cross-linking (CL)-specific enrichment of the RBPs in the interface (Fig. 2B and 2C) and provided relative quantification of the enriched RBPs (Tables S1b and S1c). Furthermore, the method revealed life cycle-selective enrichment of a set of RBPs in the parasite (Fig. 2D, Table S1d).

**FIG. 2.**
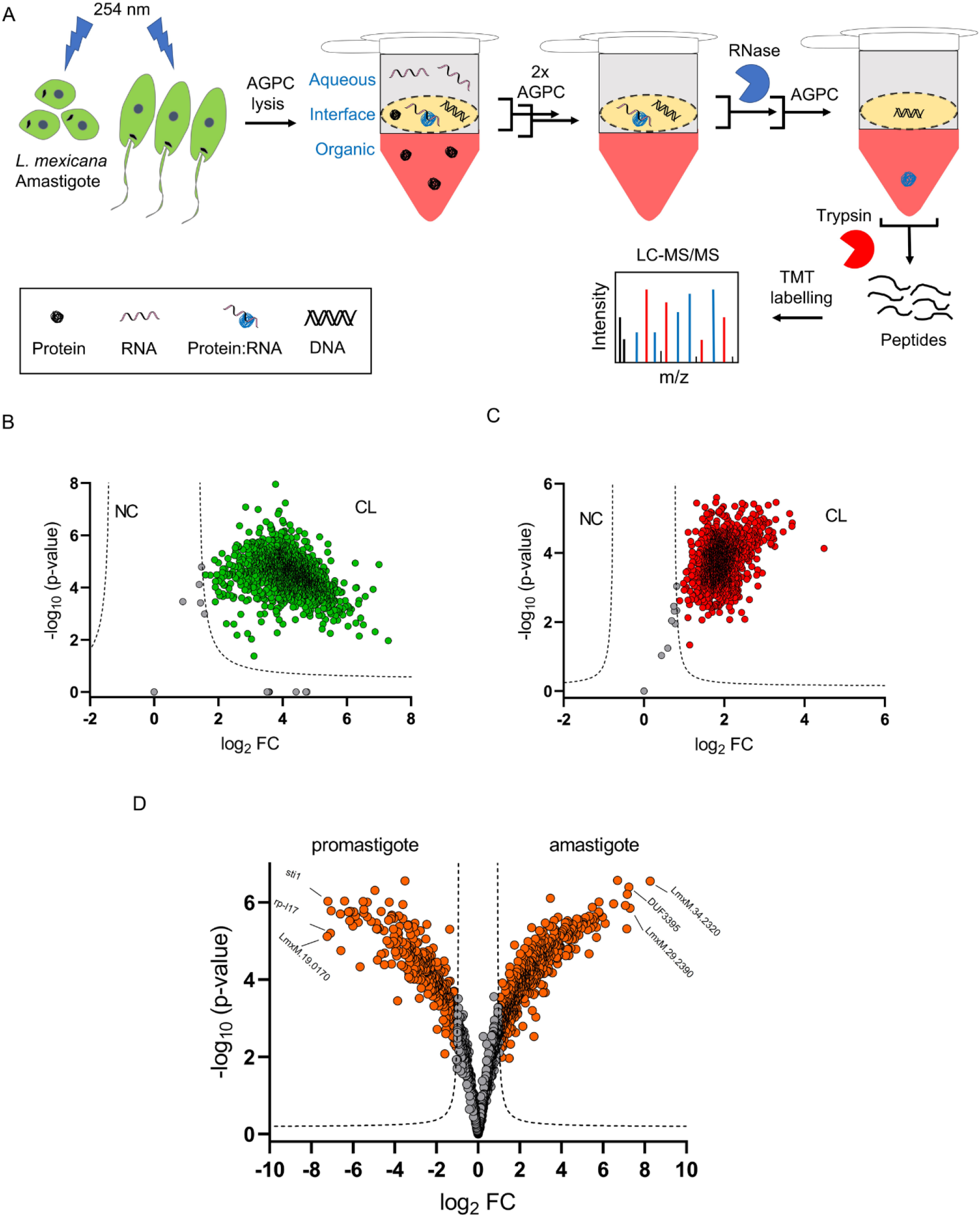
OOPS recovers RNA-binding proteins (RBPs) in *Leishmania*. (A) Schematic representation of combination of OOPS with TMT-labelling and LC-MS/MS that enables discovery and quantitation of RBPs in *L. mexicana* promastigotes and axenic amastigotes. Following UV cross-linking, cell lysis and three rounds of AGPC phase-partitioning, the third interface that is enriched with the protein-RNA adducts is collected and treated with RNase. A final AGPC phase-partitioning of the RNase-treated interface yields the previously RNA-bound proteins in the organic phase. The RBPs precipitated from the organic phase are subjected to tryptic digestion and isobaric labelling using tandem mass tags (TMT). LC-MS/MS analysis of the TMT-labelled samples provide identification of the RBPs and their fold change (FC) in the cross-linked (CL) samples compared to non-cross-linked (NC) controls. All experiments were performed in three biological replicates. Scatter plot showing CL-specific enrichment of RBPs in the OOPS interface of *L. mexicana* promastigotes (B) and axenic amastigotes (C). A modified *t*-test with permutation based FDR statistics (250 permutations, FDR = 0.001) was applied to compare three replicates of CL and NC samples in each life cycle stages. (D) Volcano plot showing differential enrichment of RBPs in the promastigote and axenic amastigote life cycle stages of *L. mexicana*. A modified *t*-test with permutation based FDR statistics (250 permutations, FDR = 0.005) was applied to compare the promastigote and amastigote groups.

A total of 2577 *L. mexicana* gene IDs in the TriTrypDB (28) were identified corresponding to the 2417 RBPs. Our total RBP data captured over 74% of mRNA-bound proteome of *L. mexicana* recently profiled using RIC (3) (Fig. 3A). Molecular Function (MF) Gene Ontology (GO)-Term analysis of the RIC data set proteins that overlapped with our OOPS data set (1232 gene IDs) showed similar RNA-specific enrichment terms as that of the additional RBPs identified in this study (Fig. 3B and 3C). However, the smaller set of 413 gene IDs in the RIC data set that have not been captured in our total RBP data set revealed no significant RNA-specific GO-enrichment Term (Fig. 3D). Thus, the OOPS method in combination with TMT labelling-based quantitative proteomic MS provided specific and comprehensive identification of RBPs in the *L. mexicana* parasites. In principle, some of the RBPs identified via the earlier RIC method (3) may bind non-poly(A) RNAs in addition to poly(A) RNAs. In contrast, the additional RBPs identified by the phase-separation method, as reported in mammalian systems recently (23), is a novel and distinct group of RNA interactors, herein designated as non-poly(A) RNA interactors in *L. mexicana*. Comparison of physicochemical properties of the poly(A)-interactors and the non-poly(A) interactors revealed similar distribution of isoelectric points versus hydrophobicity (Fig. 3E) and molecular weights (Fig. 3F). Next, we compared the interprotein domain occurrences in the two groups of RBPs, which revealed similarities and differences (Fig. 3G). In agreement with a recently reported total RBP identification in mammalian cells (23), RNA recognition motif (RRM)-containing proteins and zinc finger domains were predominantly found in the poly(A) interactome and non-poly(A) interactome respectively in the *L. mexicana*. Intriguingly, the domain with the most significant enrichment in the non-poly(A) interactome was the WD40-repeat (WDR), which often acts as scaffolds within large multiprotein complexes providing interaction sites for various biomolecules including proteins, RNA and DNA (29, 30, 31).

**FIG. 3.**
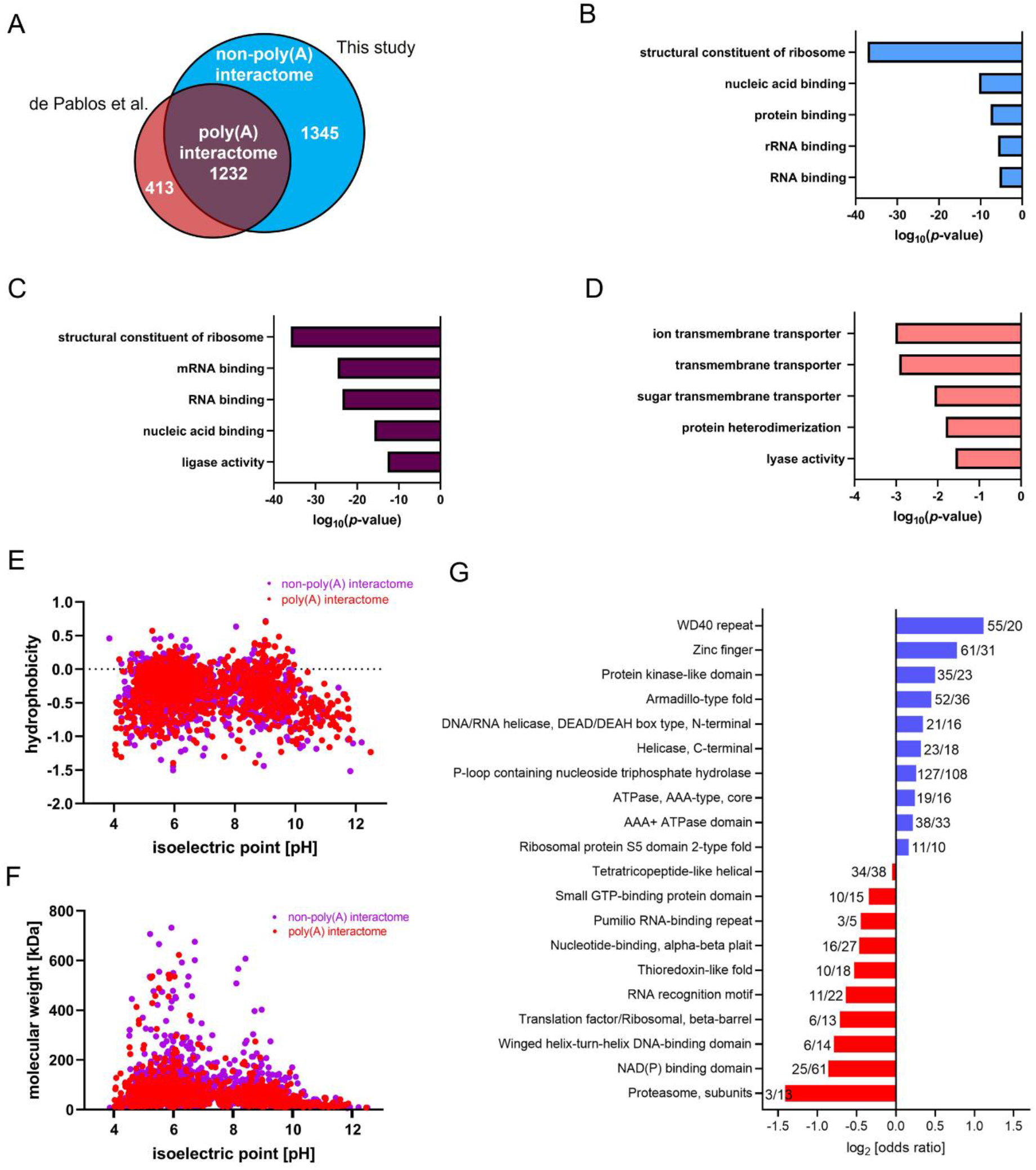
Comprehensive profiling of RBPs in *L. mexicana* by OOPS-quantitative proteomic MS combination. (A) Venn diagram showing comparison of the RBPs captured and identified by the OOPS-TMT labelling-based quantitative proteomic MS combination against those identified by the RIC-label-free quantitative proteomic MS method reported by de Pablos et al. Top five molecular function GO-Terms overrepresented in the non-poly(A) interactome (B), poly(A) interactome (C), and proteins that were exclusively present in the RIC data set (3). (D) Scatter plot comparing isoelectric points and hydrophobicity of proteins in non-poly(A) and poly(A) interactomes. (E) Scatter plot comparing isoelectric points and molecular weights of proteins in non-poly(A) and poly(A) interactomes. (F) Odd ratios of InterPro domain occurrences of non-poly(A) (blue) and poly(A) (red) interactomes.

In agreement with the extensive RNA-binding activity of proteasome proteins recently reported in mammalian cells (21, 22), a major portion of components of the *L. mexicana* proteasome complex were identified in this study (Table S2a). The proteasomal proteins were preferentially enriched in the poly(A) interactome. In contrast, eight out of twenty *L. mexicana* deubiquitinases including the recently characterised *L. mexicana* DUB2 with an elaborate RBP-interactome (32) were exclusively identified in the non-poly(A)-interactome (Table S2b), underlining the advantage of the combination of OOPS with highly sensitive TMT labelling-based quantitative proteomic MS over the RIC method in discovering RBPs with important functional roles in *Leishmania*.

### Sequencing of cross-linked and non-cross-linked RNAs in *L. mexicana*

The OOPS protocol can be applied for the large-scale discovery and validation of RBPs by combining with MS and RNA-sequencing (RNA-seq), respectively (16, 23). Having demonstrated the use of the former method in *Leishmania*, next, we compared the relative abundance of RNAs in the CL and NC samples using total RNA-seq of both promastigotes and axenic amastigotes of *L. mexicana*. Irradiation of the *Leishmania* parasites with UV-dose of 525 mJ/cm^2^ followed by sequential AGPC phase partitioning and proteinase-treatment of the final collected interface as illustrated in Fig. 1A provided the CL samples for the RNA-seq analyses. The abundance of RNA species in CL and NC samples were different (Table S3a); ncRNAs and protein coding RNAs respectively predominate the NC samples and CL samples in both promastigotes (Fig. 4A) and amastigotes (Fig. 4B). Crucially, in addition to the similar distribution of RNA species in the CL samples of both life cycle stages, the abundance of the RNAs at the interface after CL was unaffected by the size of RNAs (Fig. 4C and 4D), suggesting that similar to the mammalian system (16), the OOPS method recovered all CL *Leishmania* RNAs above 60bp without any systematic bias.

**FIG. 4.**
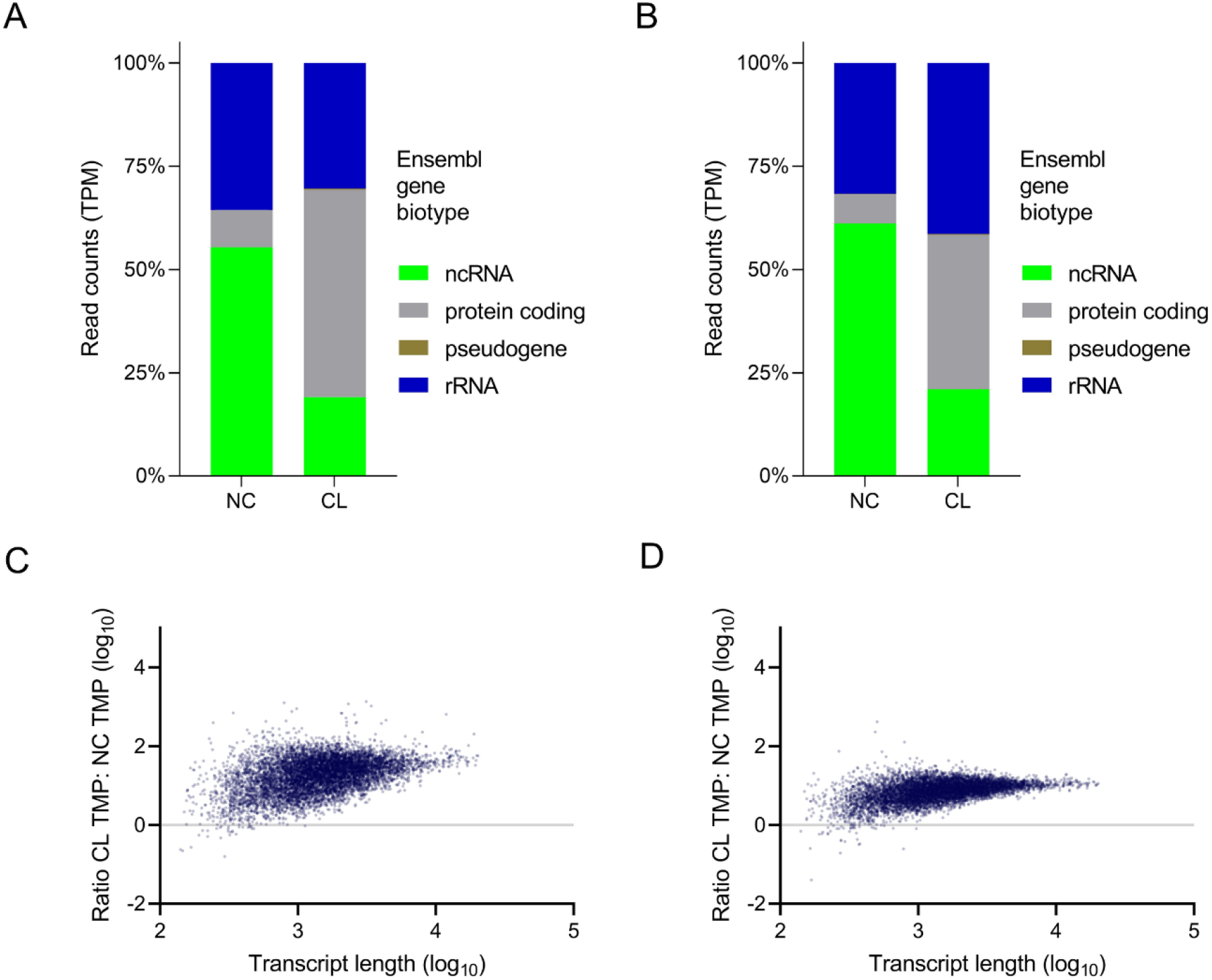
RNA-sequencing of CL and NC RNAs in *L. mexicana*. Relative proportions of RNA-seq read counts in transcripts per million (TPM) assigned to Ensembl gene biotypes for CL and NC samples in, (A) promastigotes and (B) axenic amastigotes. (F) and (G) Correlation between the ratio of CL/NC RNA and the length of the transcripts in promastigotes and axenic amastigotes respectively.

Next, we performed differential gene expression analysis of the CL versus NC samples (Fig. 5A and 5B, Tables S3b and S3c). Normalised read counts of the replicate experiments with significant (P_adj_<0.05) differential expression of transcripts in the CL versus NC samples were compared using heatmaps (Fig. 5C and 5D). Interestingly, Biological Process (BP) GO-Term analysis of the CL samples revealed a preferential enrichment of protein phosphorylation in both promastigote and axenic amastigote life cycle stages of *L. mexicana* (Fig. 5E and 5F). Of the 195 eukaryotic protein kinase genes in the *L. mexicana* genome (33), the protein-RNA-interactome captured 145 and 115 genes respectively in the promastigotes (Table S3d) and axenic amastigotes (Table S3e), suggesting a crucial role of the protein-RNA interactions in the regulation of the protein kinome of *Leishmania*.

**FIG. 5.**
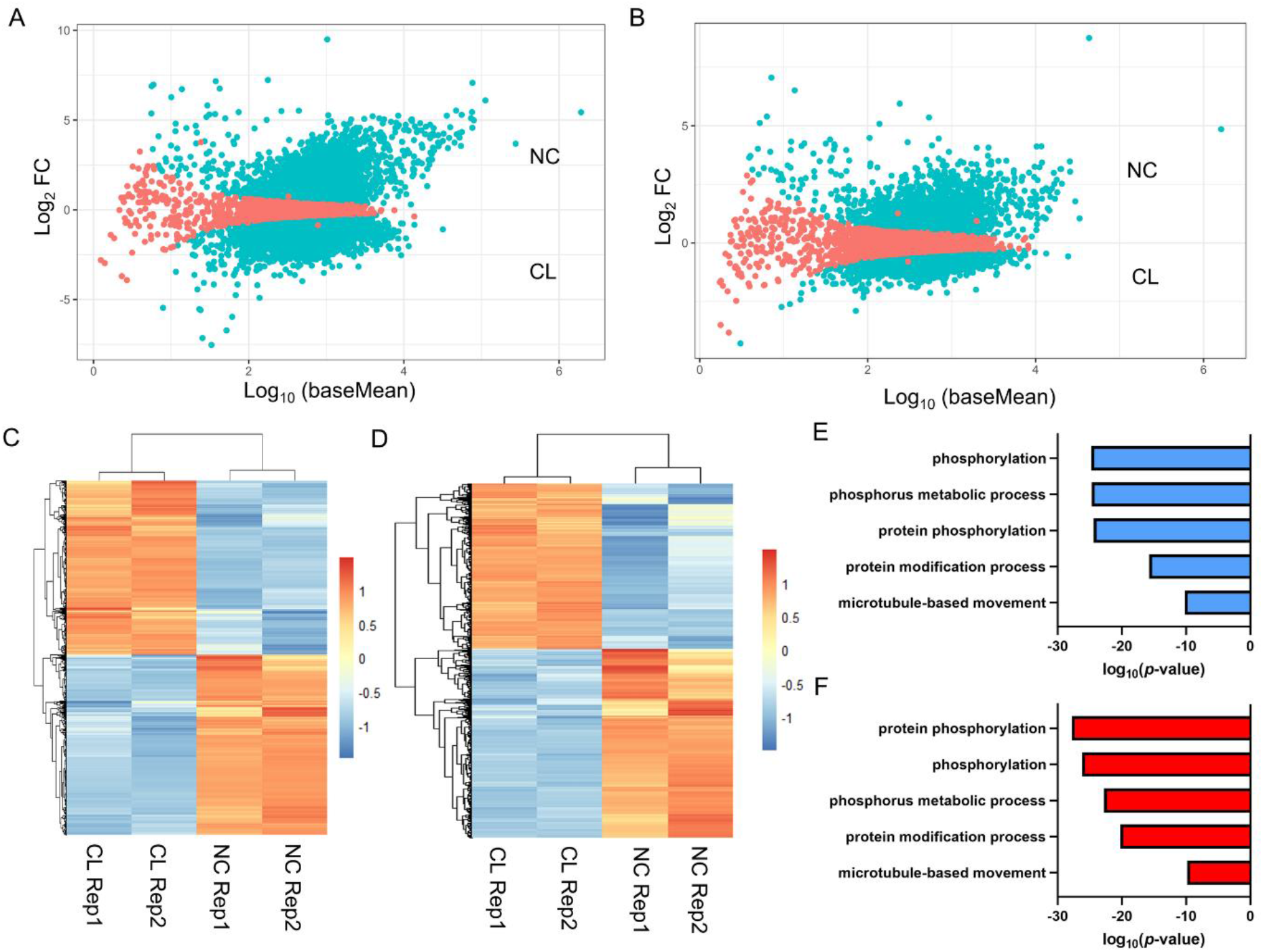
Differential gene expression analysis of RNA-seq data sets in *L. mexicana*. Scatter plot of mean of normalised read counts in log-scale (log_10_ baseMean) versus log2 fold change (log_2_ FC) for NC versus CL samples of promastigotes (A) and axenic amastigotes (B) with statistically significant hits (P_adj_<0.05) shown in blue. Heatmaps of the significant (P_adj_<0.05)DESeq2 normalised read counts of RNA-seq NC and CL replicates of promastigotes (C) and axenic amastigotes (D). Top five BP GO-Terms overrepresented in the differentially expressed (P_adj_ cut-off<0.01) CL data sets of promastigotes (E), and axenic amastigotes (F), respectively.

### Proteome-wide quantitative assessment of RBPs following Hsp90 inhibition in *L. mexicana*

Our recent study showed that the major effect of inhibition of Hsp90 in *L. mexicana* is on the parasite ribosome (34). As proteins constantly interact with RNAs in the ribosome, we asked if the Hsp90 inhibition affects the protein-RNA interactions in the parasite. To answer this, we applied the OOPS methodology in Hsp90 inhibitor treated *L. mexicana* parasites in both promastigote and axenic amastigote life cycle stages (Fig. 6A). Tryptic digests of the RNase-treated OOPS interfaces were labelled with TMT-6plex reagents such that each of the three biological replicate experiment of tanespimycin treatment versus control treatment was labelled with separate TMT duplex reagents within the TMT-6plex kit. The samples, after pooling together, were analysed by nano-LC-MS/MS. We used MS3-level spectra for accurate relative quantification between the treatments versus controls (35). A total of 2025 and 1479 proteins were identified and quantified in the promastigotes and amastigotes respectively (Tables S4a and S4c). Applying a filter of minimum 2 unique peptides and 3 valid values across the three replicates, a total of 891 and 496 statistically significant high-confidence tanespimycin-affected RBPs were identified in the promastigotes and amastigotes respectively (Tables S4b and S4d).

**FIG. 6.**
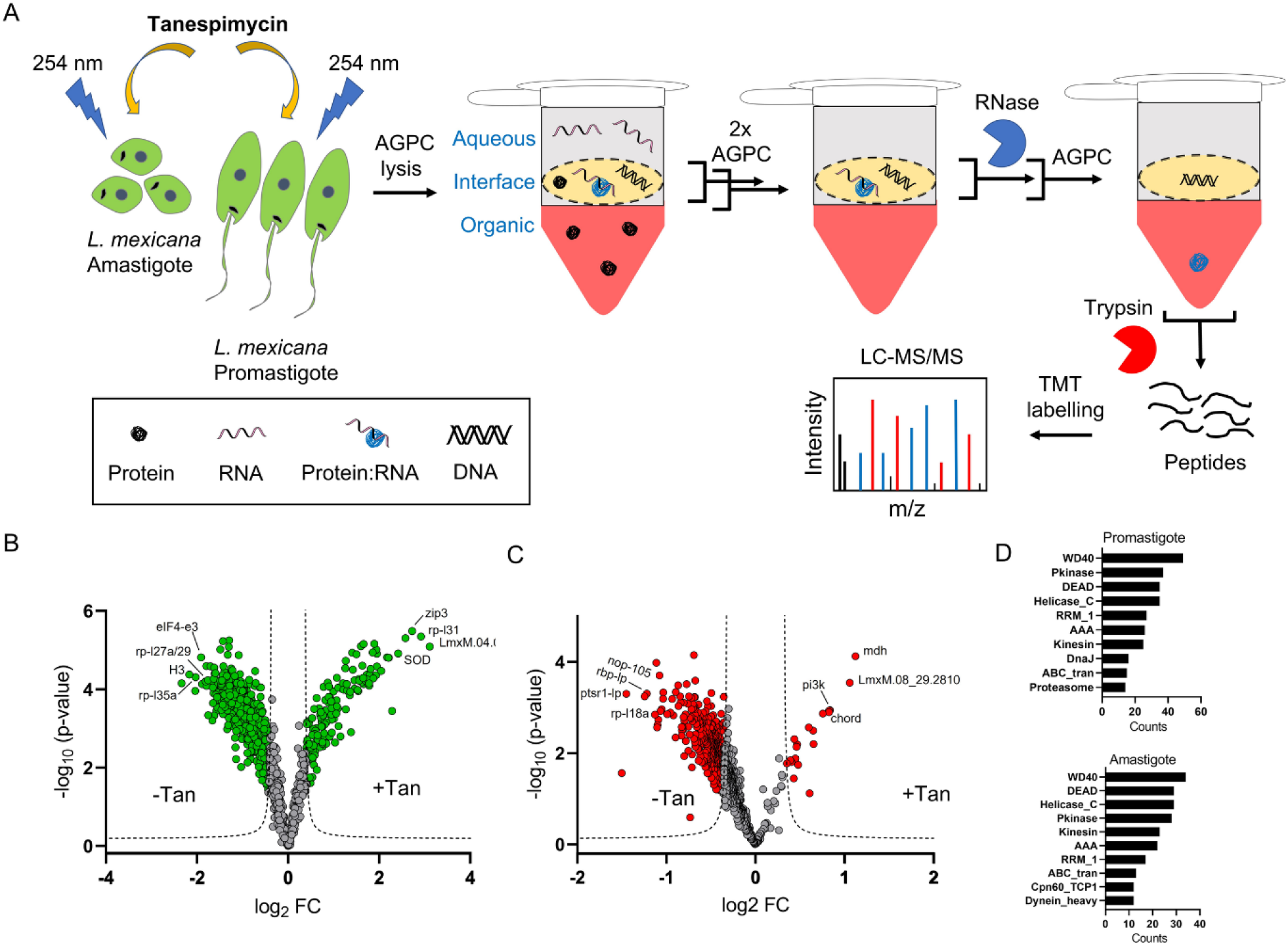
Hsp90 inhibition affects protein-RNA interactions in *Leishmania*. (A) Schematic representation of combination of Hsp90 inhibition with tanespimycin treatment and the OOPS-quantitative proteomic MS method for proteome-wide quantitative assessment of the effect of the inhibition on protein-RNA interactions in both promastigote and axenic amastigote life cycle stages of *L. mexicana*. For the Hsp90 inhibition, the parasites in three independent replicates were treated with 1 µM tanespimycin or vehicle (DMSO) for 16 h prior to the OOPS protocol. Volcano plots showing differential regulation of RBPs upon tanespimycin treatment (+Tan) with respect to vehicle treatment (-Tan) in promastigotes (B) and axenic amastigotes (C). A modified *t*-test with permutation based FDR statistics (250 permutations, FDR = 0.05) was applied to compare the +Tan and -Tan groups in both life cycle stages. (D) Top-10 Pfam domain occurrences in tanespimycin-affected RBPs in promastigotes and axenic amastigotes.

Whilst tanespimycin caused an increase in the RNA-binding of a small set of RBPs, the RNA-binding of the majority of RBPs were negatively affected in both life cycle stages of *L. mexicana* (Fig. 6B and 6C). Interestingly, different set of RBPs were observed among the mostly affected in the promastigotes and amastigotes, contrasting the molecular signatures of the Hsp90 inhibition on the RBPs of the two life cycle stages (Table S4e). The effect of tanespimycin on the RNA binding of proteins was largely independent of the relative levels of the RBPs quantified between the two life cycle stages (Table S4e, Supplementary Fig. S2A, S2B). Despite these differences the isoelectric point, hydrophobicity, and molecular weight of the affected proteins in both life cycle stages showed similar patterns (Supplementary Fig. S2C to S2D). Additionally, a comparison of domain occurrences revealed commonalities between the affected RBPs of the two stages (Fig. 6D). Similarly, in both life cycle stages, GO analyses of the affected proteins revealed structural constituent of ribosome (P values 1.14e^-37^ for promastigotes and 4.50e^-42^ for amastigotes), ribosome (P values 4.00e^-23^ for promastigotes and 1.43e^-23^ for amastigotes), and translation (P values 2.94e^-47^ for promastigotes and 4.08e^-50^ for amastigotes) as extremely enriched MF, Cellular Component (CC) and BP GO-Terms respectively (Table S4f), agreeing with our recent finding that the major effect of inhibition of Hsp90 in *L. mexicana* is on the parasite ribosome (34). However, the downstream molecular effects of the inhibition beyond protein expression remained unknown. Our combination of the Hsp90 inhibition with the OOPS-based total RBP discovery in *L. mexicana* revealed that the Hsp90 inhibition causes large-scale disruption of the RNA binding of several ribosomal protein (RPs), proteasomal proteins and translation factors in both promastigote and amastigote life cycle stages of the parasite (Tables S4b and S4d).

Next, we intersected the *L. mexicana* promastigote RBP-tanespimycin treatment data set with our recently reported quantitative proteomic data set of effect of tanespimycin treatment on protein synthesis in the parasite (34). Out of 173 proteins quantified between the two data sets, we observed positive and negative correlation between protein synthesis and RNA binding on 101 and 72 proteins respectively (Table S5). Among these, the RNA binding of 49 and 15 structural constituent of ribosome proteins showed positive and negative correlation respectively with their expression levels (Table S5). Whilst the RNA binding of most RPs was decreased upon tanespimycin treatment, striking increase in the RNA binding independent of the change in the expression was observed on four RPs; 60S RPL31 (LmxM.34.3280, LmxM.34.3290), 40S RPS7 (LmxM.01.0410), 40S RPS11 (LmxM.21.1550) and 60S RPL34 (LmxM.36.3740), suggesting that specific RPs bind RNA differentially and the Hsp90 inhibition differentially disrupts specific RNA-protein interactions.

## DISCUSSION

Protein-RNA interactions regulate a myriad of processes that are crucial for the cell to maintain homeostasis. Although mRNAs constitute only 2 to 5% of the total transcripts in eukaryotes, until very recently the identification of RBPs was limited to mRNA-binding proteins. This was because of the limitation of the RIC technique, which relies on an affinity interaction between the poly(A) tails in mRNAs and oligo(dT) beads. In addition to missing out the protein interactors of the vast majority of transcripts in the eukaryotic cell, RIC cannot be applied in bacteria and many archaea as they lack poly(A) tails in their transcripts. Noncoding RNAs (ncRNAs) participate in numerous biological functions (14, 15). The highly abundant ncRNAs, the rRNAs, serve not only as the major structural constituents of ribosome, but also associate with RPs and facilitate peptidyl transfer reaction in the protein synthesis (36). The tRNAs also play a pivotal role in protein synthesis by serving as adaptor molecules for amino acids (37). The functional roles of other ncRNAs including long noncoding RNAs (lncRNAs), small nuclear RNAs (snRNAs) and small nucleolar RNAs (snoRNAs) is only beginning to unravel. As most coding and noncoding RNAs function as RNPs, identification of the total RBPs is of paramount importance for deciphering the underlying biological processes they regulate.

We combined the OOPS method with TMT labelling based quantitative proteomic MS for robust transcriptome-wide discovery of coding and noncoding RBPs in both promastigote and amastigote life cycle stages of *L. mexicana* parasites. With about two times the number of RBPs identified compared to a recently reported large-scale study that employed RIC and label-free quantitative proteomics (3), our study provides the most comprehensive data set of RBPs in *L. mexicana*. Importantly, our study unravelled an extensive set of non-poly(A) interactors in the parasite, which the RIC method could not capture. Interprotein domain analysis of the non-poly(A) interactors revealed a preferential enrichment of WDR domains. Although the functional roles of the WDR domains are only beginning to emerge, they often serve as crucial units of large multiprotein complexes mediating diverse functions including RNA processing, ubiquitin signalling, protein degradation, sensing of DNA damage, DNA repair, control of cell division, chromatin organisation and epigenetic regulation of gene expression (29, 30, 31). Structural basis of sequence-specific RNA binding by WDR domains have been reported by both crystallographic studies (38, 39) and high-resolution cryo-electron microscopy studies (40,41, 42, 43, 44). In humans many proteins that harbour WDR domains are disease-associated (30), and the WDR domains of two proteins, WDR5 and EED, have been specifically targeted by potent and cell-active drug-like small molecules (45, 46, 47). These small molecule chemical probes have revealed high structural diversity of the binding pockets in the WDR domains, suggesting druggability of the ubiquitous protein domain. The prevalence of the WDR domains among the novel class of non-poly(A) RNA interactors of *L. mexicana* revealed by our study therefore will fuel both functional studies and investigation of their druggability scope in the *Leishmania* spp. parasite. Studies in this direction are currently in progress in our group.

In agreement with the RIC data set (3), we identified an extensive set of *L. mexicana* proteasomal proteins as RBPs. Whilst the biological functions of the RNA-binding activity of the proteasome proteins remain elusive the 20S core of proteasome is reported to possess RNase activity (48, 49). Also, several proteins in the 19S regulatory particle of proteasome participate in functions independent of the proteasome complex (50). However, in contrast to the RIC data set that identified only a limited number of ubiquitin-associated proteins, our OOPS-TMT-6plex quantitative proteomics method identified an elaborate list of ubiquitin-associated proteins, including 10 out of 20 deubiquitinases in the parasite, suggesting hitherto unidentified and potentially widespread role of deubiquitinases in the processing of RNAs in the *Leishmania* parasite. DUB2, the top RNA-binding deubiquitinase detected in our OOPS data set, has been recently identified to be an essential protein required for life cycle transition in the *L. mexicana* (32). The DUB2 interactome profiling in *L. mexicana* has revealed its physical association with several RBPs involved in transcription, chromatin dynamics, mRNA splicing and RNA capping (32). Also, RNA-binding ubiquitin ligases have been suggested to act as key links between posttranscriptional regulation and the ubiquitin system (51).

The identification of many proteasomal proteins, deubiquitinases, ubiquitin conjugating enzymes and ligases suggest potential functional roles of the protein-RNA interactions in crucial cellular processes such as protein degradation by the ubiquitin-proteasome system (UPS) and proteasomal degradation. Targeting of the proteasome in *Leishmania* by small molecules has been recently reported as a promising strategy for the development of therapeutic agents against leishmaniasis (52, 53, 54). The widespread RNA-binding property identified in the proteasome proteins suggests functional relevance of the protein-RNA interactions and demands further investigation.

Transcriptome-profiling by total RNA-seq of the NC and CL samples revealed similar relative proportions of Ensembl gene biotypes in the RNA-seq reads of the CL samples of promastigotes and amastigotes, contrasting the CL samples from the NC samples. The higher relative proportion of protein coding genes in the CL samples underscores the importance of protein-RNA interactions in the regulation of gene expression in *Leishmania*. Crucially, the CL samples also contain a substantial proportion of the ncRNAs, suggesting potential functional roles of the protein-binding ncRNAs in the posttranscriptional regulation of gene expression in the protozoan parasite. The enrichment of protein phosphorylation BP GO-Term and a substantial portion of the *L. mexicana* protein kinase genome in the CL samples, revealed by differential gene expression analysis, suggests potential role of the RBPs in the large-scale regulation of the protein kinome of *Leishmania*. The *L. mexicana* possess an extensive set of eukaryotic protein kinases and a recent study has shown that many of the protein kinases act as regulators of life cycle differentiation and survival in the parasite (33).

The combination of Hsp90 inhibition with the OOPS-TMT-6plex based total RBP identification showed that whilst largely different set of RBPs were affected by the inhibitor in the promastigotes and amastigotes, the overall cellular effect of the inhibition as revealed by the GO analyses of the affected RBPs were strikingly similar in both life cycle stages. This suggests that the parasite relies on different repertoire of protein-RNA interactions to cope up with the Hsp90 inhibition stress in the two different life cycle stages, but nevertheless accomplishes similar end results, underscoring the extraordinary ability of *Leishmania* to adapt to stress conditions. We observed RNA binding of RPs, proteasomal proteins, and translation factors amongst the mostly affected, suggesting protein-RNA interactions playing crucial roles in the downstream effect of the Hsp90 inhibition on protein synthesis and degradation pathways in the parasite. This study thus provides a wealth of insights for further studies to illuminate the molecular mechanisms of Hsp inhibition in the *Leishmania* spp. parasites.

## MATERIALS AND METHODS

### *Leishmania* cell culture

*L. mexicana* strain M379 (MNYC/BC/62/M379) promastigotes were grown in T-75 flasks at 26 °C in Schneider’s insect medium (Sigma-Aldrich) supplemented with 0.4 g/L NaHCO_3_, 0.6 g/L anhydrous CaCl_2_, and 10% heat-inactivated foetal bovine serum (FBS) (pH 7.0). Axenic amastigote cultures were generated from procyclic promastigotes using changes in the pH and temperature of the culture medium. Briefly, *L. mexicana* promastigotes in log stage were seeded at 5 × 10^5^ parasites/mL in 30 mL pH 7.0 Schneider’s insect medium supplemented with 15% heat-inactivated FBS and incubated at 26 °C. On day 3, the procyclic promastigotes were transferred to 60 mL of pH 5.5 Schneider’s insect medium supplemented with 20% heat-inactivated FBS, seeded at 5 × 10^5^ parasites/mL and incubation continued at 26 °C. On day 7, the metacyclic parasites were transferred to 60 mL of pH 5.5 Schneider’s insect medium supplemented with 20% heat-inactivated FBS, seeded at 5 × 10^5^ parasites/mL and incubated at 32 °C. On day 11-12, the parasites were completely differentiated into amastigote stage. The growth and morphology of parasites were observed under an optical microscope and parasite numbers in cultures were measured using a haemocytometer.

### OOPS in *L. Mexicana*

Mid-log phase promastigotes and axenic amastigotes grown in T-75 flasks (30 mL culture medium; 5 × 10^6^ parasites/mL) were pelleted by centrifugation (3 min at 900g, room temperature) and washed once with PBS. In non-cross-linked controls, 100 µL PBS was added to the pellets, the parasites were passed through 29G needle (5 times) and subjected to acidic guanidinium thiocyanate-phenol-chloroform (AGPC) biphasic extraction as reported earlier for mammalian and bacterial cells (16). Briefly, one mL Trizol reagent was added to each sample and homogenised by pipetting several times followed by vortexing at maximum speed for 15 seconds. For biphasic extraction, 200 µL chloroform (Fisher Scientific) was added, vortexed and the samples were centrifuged at 12,000g for 15 min at 4 °C. In cross-linked samples, parasites were resuspended in PBS in 6-well plates and cross-linked in solution at 254 nm with UV dosages varying from 75 mJ/cm^2^ to 650 mJ/cm^2^ (CL-3000 Ultraviolet Crosslinker; UVP). For quantitative proteomic experiments, the parasites were irradiated with UV dosage of 525 mJ/cm^2^. The cross-linked parasites were pelleted, and supernatant removed by pipetting leaving approximately 100 µL PBS. The parasites were passed through 29G needle (5 times), Trizol reagent was added and homogenised by pipetting several times followed by vortexing at maximum speed for 15 seconds as described above. The homogenised lysates were incubated at room temperature (RT) for 5 min to dissociate unstable RNA-protein interactions, and AGPC biphasic extraction was performed as described above. The upper aqueous phase containing non-cross-linked RNAs was transferred to a new tube and RNA isolated by standard phenol/chloroform extraction (Thermo Fisher Scientific). The lower organic phase containing non-cross-linked proteins was precipitated by addition of nine volumes of methanol. The interface containing the RNA-protein adducts was transferred to a new tube and subjected to two additional rounds of AGPC biphasic extraction. The interface of the final AGPC phase separation cycle was precipitated by addition of nine volumes of methanol and pelleted by centrifugation at 14,000g for 10 min at RT.

### RNA extraction and quantification

The precipitated interfaces were incubated with 20 U of proteinase K (Thermo Fisher Scientific) in a protein digestion buffer (30 mM Tris HCl, pH 8:00, 10 mM EDTA) at 50 °C for 2 h. Samples were cooled to RT and released RNA was purified by standard phenol/chloroform extraction (Thermo Fisher Scientific) following manufacturer’s instructions. RNA purity was assessed by Nanodrop (DeNovix). Samples with a 260/230 absorbance ratio below 2 and 260/280 absorbance ratio below 1.9 were discarded. RNA concentration was determined using QuantiFluor RNA system (Promega) in BioTek Synergy H4 plate reader in 96-well black flat-bottom plates following manufacturer’s instructions.

### RNA sequencing

Total non-crosslinked (NC) RNA and protein-cross-linked (CL) RNA from promastigote and axenic amastigote life cycle stages of *L. mexicana* parasites in biological duplicates were purified using standard Trizol extraction and OOPS, respectively. The CL samples were treated with 20 U of proteinase K in a protein digestion buffer (30 mM Tris HCl, pH 8:00, 10 mM EDTA) at 50 °C for 2 h and the released RNA was purified by a subsequent Trizol extraction. All RNA samples were treated with Turbo DNase (Thermo Fisher Scientific). RNA integrity was evaluated using Agilent 2100 Bioanalyzer system and sample purity was further assessed using agarose gel electrophoresis. Ribosomal RNA (rRNA) was removed by NEBNext rRNA Depletion Kit (New England BioLabs) according to manufacturer instructions, and sequencing libraries were generated using the rRNA-depleted RNA samples using NEBNext Ultra Directional RNA Library Prep Kit for Illumina (New England BioLabs). Briefly, after fragmentation, the first strand cDNA was synthesised using random hexamer primers. Then the second strand cDNA was synthesised and dUTPs were replaced with dTTPs in the reaction buffer. The directional library was ready after end repair, A-tailing, adapter ligation, size selection, USER enzyme digestion, amplification, and purification. The library was checked with Qubit real-time PCR for quantification and bioanalyzer for size distribution detection. All libraries were sequenced in parallel on a NovaSeq 6000 PE150 (Illumina).

### RNA-seq data processing and bioinformatics

*Leishmania mexicana* MHOM/GT/2001/U1103 genome sequences, transcriptome sequences and general feature format (gff) file were downloaded from TriTrypDB release 54 (28). Counts of reads per gene were obtained using STAR aligner (55). Relative abundance of transcripts in units of transcripts per million (TPM) was obtained using Salmon (56) with default settings. GraphPad Prism version 9.2.0 (www.graphpad.com) was used for generating scatter plots, and for performing correlation analyses. Differential gene expression analyses were performed in R (version 4.1.1) using the Bioconductor DESeq2 package (57) and plots were generated using the R package ggplot2.

### Proteomic sample preparation and TMT labelling

Protein samples were resuspended in 200 µL of 100 mM triethylammonium bicarbonate (TEAB) (Sigma-Aldrich), reduced with 20 mM dithiothreitol (DTT) (Sigma-Aldrich) at RT for 60 min and alkylated with 40 mM iodoacetamide (Sigma-Aldrich) at RT in the dark for 60 min. Samples were digested overnight at 37 °C with 5 µg sequencing-grade modified trypsin (Promega). The samples were then acidified with trifluoroacetic acid (TFA) (0.1% v/v final concentration; Sigma-Aldrich), centrifuged at 16,000g for 10 min and the supernatant was collected. The tryptic peptides were then desalted on C-18 Sep-Pak Classic cartridges (Waters; WAT051910) following manufacturer’s instructions. The peptides were evaporated to complete dryness under a vacuum and stored at -80 °C until required.

TMT labelling of the desalted tryptic peptides was carried out using TMTsixplex isobaric label reagent set (Thermo Fisher Scientific). The peptides of each experimental condition were dissolved in 100 µL of 100 mM TEAB and treated with an RT equilibrated and freshly dissolved unique TMT label reagent in 41 µL anhydrous acetonitrile. The labelling reactions were run for 1 h at RT, following which 10 µL of 5% solution of hydroxylamine was added and incubated for 15 min at RT to quench the reactions. The six samples of three replicates of each life cycle stage were then combined together and concentrated to complete dryness under a vacuum. The samples were then redissolved in 0.1% TFA, desalted and cleaned-up using Pierce Peptide Desalting Spin Columns (Thermo Fisher Scientific) following manufacturer’s instructions. The samples were dried on a SpeedVac and stored at -80 °C until required.

### LC-MS/MS

The LC-MS/MS analysis of TMT-labelled peptides were performed on an Orbitrap Fusion Lumos mass spectrometer (Thermo Fisher Scientific) coupled with a Thermo Scientific Ultimate 3000 RSLCnano UHPLC system (Thermo Fisher Scientific). The peptides dissolved in 0.1% formic acid (FA) were first loaded onto an Acclaim PepMap 100 C18 trap column (5 µm particle size, 100 µm id X 20 mm, TF164564) heated to 45 °C using 0.1% FA/H_2_O with a flow rate of 10 µL/min, then separated on an Acclaim PepMap 100 NanoViper C18 column (2 µm particle size, 75 µm id X 50 cm, TF164942) with a 5% to 38% ACN gradient in 0.1% FA over 80 min (*L. mexicana* amastigote and promastigote OOPS-TMT-6plex MS2 runs) or over 120 min (*L. mexicana* amastigote and promastigote tanespimycin-OOPS-TMT-6plex MS3 runs) at a flow rate of 300 nL/min. The full MS spectra (*m/z* 375 to 1,500) were acquired in Orbitrap at 120,000 resolution with an AGC target value of 4e^5^ for a maximum injection time of 50 ms. High-resolution HCD MS2 and MS3 spectra were generated in positive ion mode using a normalised collision energy of 36% and 65% respectively within a 0.7 *m/z* isolation window using quadrupole isolation. The AGC target value was set to 10e^4^, and the dynamic exclusion was set to 45 s. The MS2 spectra were acquired in Orbitrap with a maximum injection time of 86 ms at a resolution of 50,000 with an instrument determined scan range beginning at *m/z* 100. The MS3 spectra were acquired in Orbitrap with a maximum injection time of 100 ms at a resolution of 50,000. The scan range of MS3 was *m/z* 100 to 500. To ensure quality peptide fragmentation a number of filters were utilised, including peptide monoisotopic precursor selection, minimum intensity exclusion of 10e^3^ and exclusion of precursor ions with unassigned charge state as well as charge state of +1 or superior to +7 from fragmentation selection. To prevent repeat sampling, a dynamic exclusion with exclusion count of 1, exclusion duration of 45 ms, mass tolerance window of +/-7 ppm and isotope exclusion were used.

### MS spectra processing and protein identification and quantification

Raw data were processed using MaxQuant software version 1.5.3.30 (58) with integrated Andromeda database search engine (59). The MS/MS spectra were queried against *L. mexicana* sequences from UniProt KB (8,524 sequences). The following search parameters were used: reporter ion MS2 with multiplicity 6plex TMT for the amastigote and promastigote OOPS-TMT-6plex MS2 runs and reporter ion MS3 with multiplicity 6plex TMT for the amastigote and promastigote tanespimycin treatment-OOPS-TMT-6plex MS3 runs, trypsin digestion with maximum 2 missed cleavages, carbamidomethylation of cysteine as a fixed modification, oxidation of methionine and acetylation of protein N-termini as variable modifications, minimum peptide length of 6, maximum number of modifications per peptide set at 5, and protein false discovery rate (FDR) 0.01. Appropriate correction factors for the individual TMT channels for both lysine side-chain labelling and peptide N-terminal labelling as per the TMT-6plex kits used (Thermo Fisher Scientific) were configured into the database search. The proteinGroups.txt files from the MaxQuant search outputs were processed using Perseus software version 1.6.2.3 (60). Sequences only identified by site, reverse sequences, and potential contaminants were filtered out. A requirement of six non-zero valid value were set across the six replicates within the OOPS-TMT-6plex MS2 experiments. For the tanespimycin treatment OOPS-TMT-6plex MS3 runs, a requirement of three non-zero valid value were set across the 3 replicates of the amastigote and promastigote samples separately. Proteins identified with fewer than 2 unique peptides were discarded and a modified *t-*test with permutation-based FDR statistics was applied (250 permutations) to compare the CL and NC groups, amastigote and promastigote samples, and tanespimycin treated and non-treated groups.

### Bioinformatic analysis

GO Terms (Molecular Function, Biological Process, and Cellular Component) of the RBP data sets were derived from TriTrypDB (tritrypdb.org) (28). InterPro domain occurrences in the poly(A) and non-poly(A) interactomes were derived from bioinformatics analysis of the protein IDs using the Database for Annotation, Visualization and Integrated Discovery (DAVID) v6.8 bioinformatics resources (61). Grand average hydrophathicity values of the RBPs were calculated using the ExPASy (62) tool ProtParam. Isoelectric points and molecular weights were computed using the ExPASy tool Compute pI/Mw.

### Data availability

All raw mass spectrometry proteomics data have been deposited to the ProteomeXchange Consortium via the PRIDE partner repository with the dataset identifiers PXD028092. All sequencing data can be accessed through the NCBI Gene Expression Omnibus (GEO) genomic data repository with the accession code GSE188357.

## SUPPLEMENTAL MATERIAL

**FIG. S1**, TIF file

**FIG. S2**, TIF file

**TABLE S1**, XLSX file

**TABLE S2**, XLSX file

**TABLE S3**, XLSX file

**TABLE S4**, XLSX file

**TABLE S5**, XLSX file

## ACKNOWLEDGEMENTS

This work of K. K, B. S. M, and P. W. D was supported by funding from MRC-Global Challenges Research Fund-Neglected Tropical Diseases, Grant number: MR/P027989/1A. The work of T. I. R and J. C was funded by the CRUK Centre grant with reference number: C309/A25144.

## SUPPLEMENTAL MATERIAL LEGEND

**FIG. S1** Absolute quantification of the total recovered RNA (PBR + free RNA) in *L. mexicana* promastigotes and axenic amastigotes. Data shown as mean +/-SD of three independent experiments.

**FIG. S2** Tanespimycin affects the RBPs of *L. mexicana*. Volcano plots showing differential enrichment of RBPs in the promastigote and axenic amastigote life cycle stages of *L. mexicana* with vehicle (DMSO) treatment (A) and with tanespimycin (1 µM for 16 h) treatment (B). A modified *t*-test with permutation based FDR statistics (250 permutations, FDR = 0.05) was applied to compare the +Tan and -Tan groups. (C) and (D), Scatter plots comparing isoelectric points and hydrophobicity of proteins in promastigotes and amastigotes respectively. (E) and (F) Scatter plots comparing isoelectric points and molecular weight of proteins in promastigotes and amastigotes respectively.

**TABLE S1** RBPs of *L. mexicana* profiled and quantified by OOPS-TMTsixplex-LC-MS/MS.

a) Total RBPs identified with two or more unique peptides and a minimum of 6 valid reporter ion quantification across 12 samples. b) and c) Total RBPs identified with two or more unique peptides in promastigotes and amastigotes respectively. The fold change (FC) of CL samples versus NC samples in log2 scale and statistical significance of the reporter intensity quantifications across three independent replicates of CL versus NC samples as negative log of *t*-test P-values are also presented d) RBPs of promastigotes compared with those of amastigotes across the three CL replicates of both life cycle stages.

**TABLE S2** *L. mexicana* poly(A) and non-poly(A) interactomes.

**TABLE S3** RNA-seq results. a) read counts of the NC and CL replicate samples of *L. mexicana* promastigotes (Proma) and axenic amastigotes (Ama). b) and c) differential gene expression analysis by DESeq2 R package of NC versus CL samples of promastigotes and axenic amastigotes respectively. d) and e) protein kinase genes identified in the CL transcripts of *L. mexicana* promastigotes and axenic amastigotes respectively.

**TABLE S4** Hsp90 inhibition by tanespimycin affects RNA-protein interactions in *L. mexicana*. a) and c) total RBPs profiled with tanespimycin (+Tan) and without tanespimycin (-Tan) treatment using the OOPS-TMTsixplex-LC-MS/MS in promastigotes and axenic amastigotes respectively. b) and d) High-confidence tanespimycin-affected RBPs of promastigotes and axenic amastigotes respectively. Ribosomal proteins, proteasomal proteins and translation factors are highlighted in red, blue, and green respectively. e) Tanespimycin-affected RBPs compared between the amastigote and promastigote groups. f) Top-10 MF, CC and BP GO-Terms enriched in the tanespimycin-affected RBPs of *L. mexicana*.

**TABLE S5** Effect of tanespimycin on the RBPs of *L. mexicana* promastigotes compared with the effect on the synthesis of the nascent proteins revealed by BONCAT-iTRAQ quantitative proteomic MS (ref. 32). An increase (positive value of log2 FC) or decrease (negative value of log2 FC) of RNA-binding upon tanespimycin-treatment quantified by the MS3 dataset of the OOPS-TMTsixplex-LC/MS/MS is compared with the change in abundance of the nascent proteins revealed by the BONCAT-iTRAQ quantitative proteomic MS data set. Positively and negatively correlated RBPs are highlighted in blue and orange respectively.

