## Supplementary figures and images for "Transcriptome-wide identification of coding and noncoding RNA-binding proteins defines the comprehensive RNA interactome of *Leishmania mexicana*"

### Figure S1

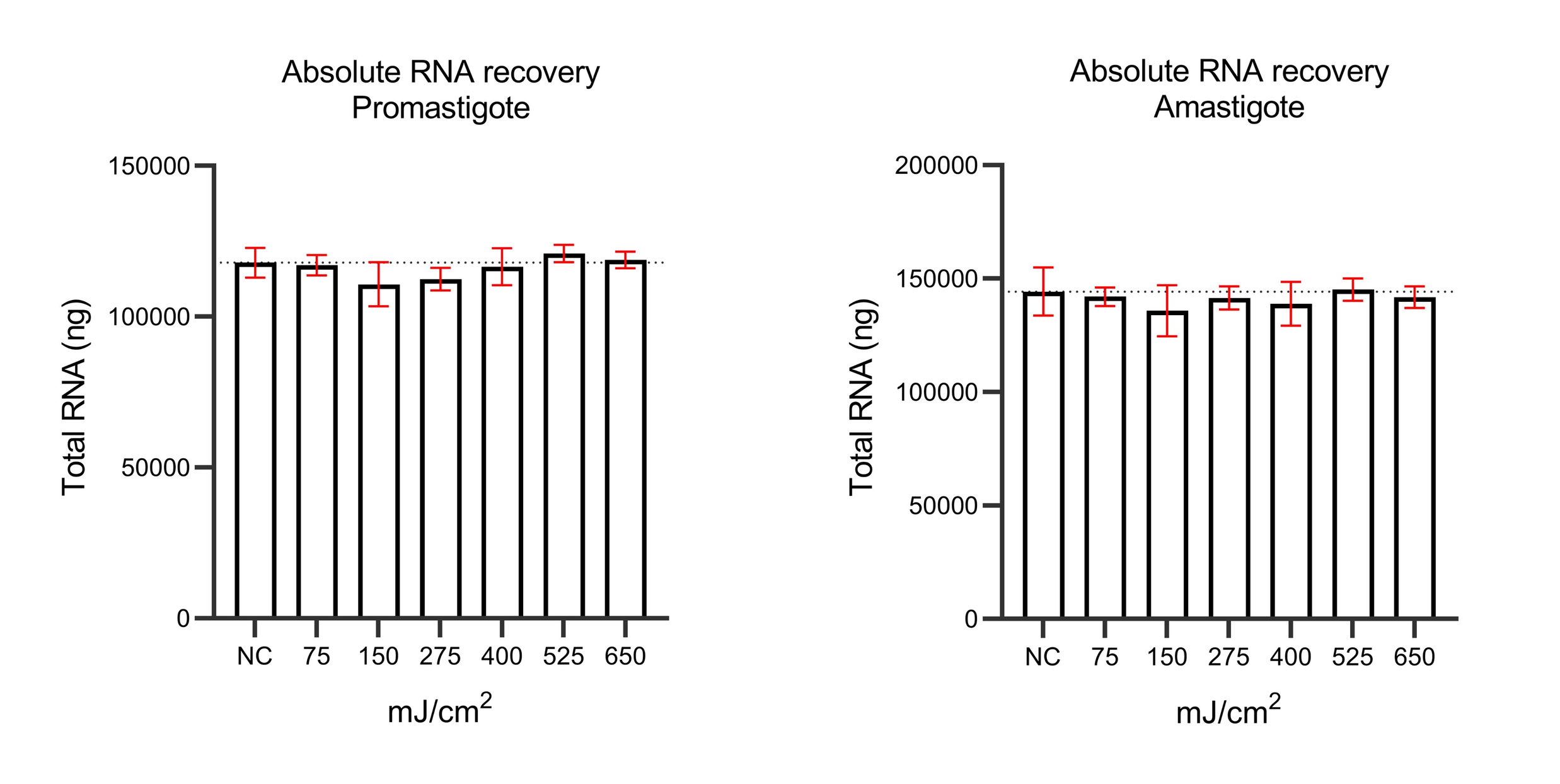

### Figure S2

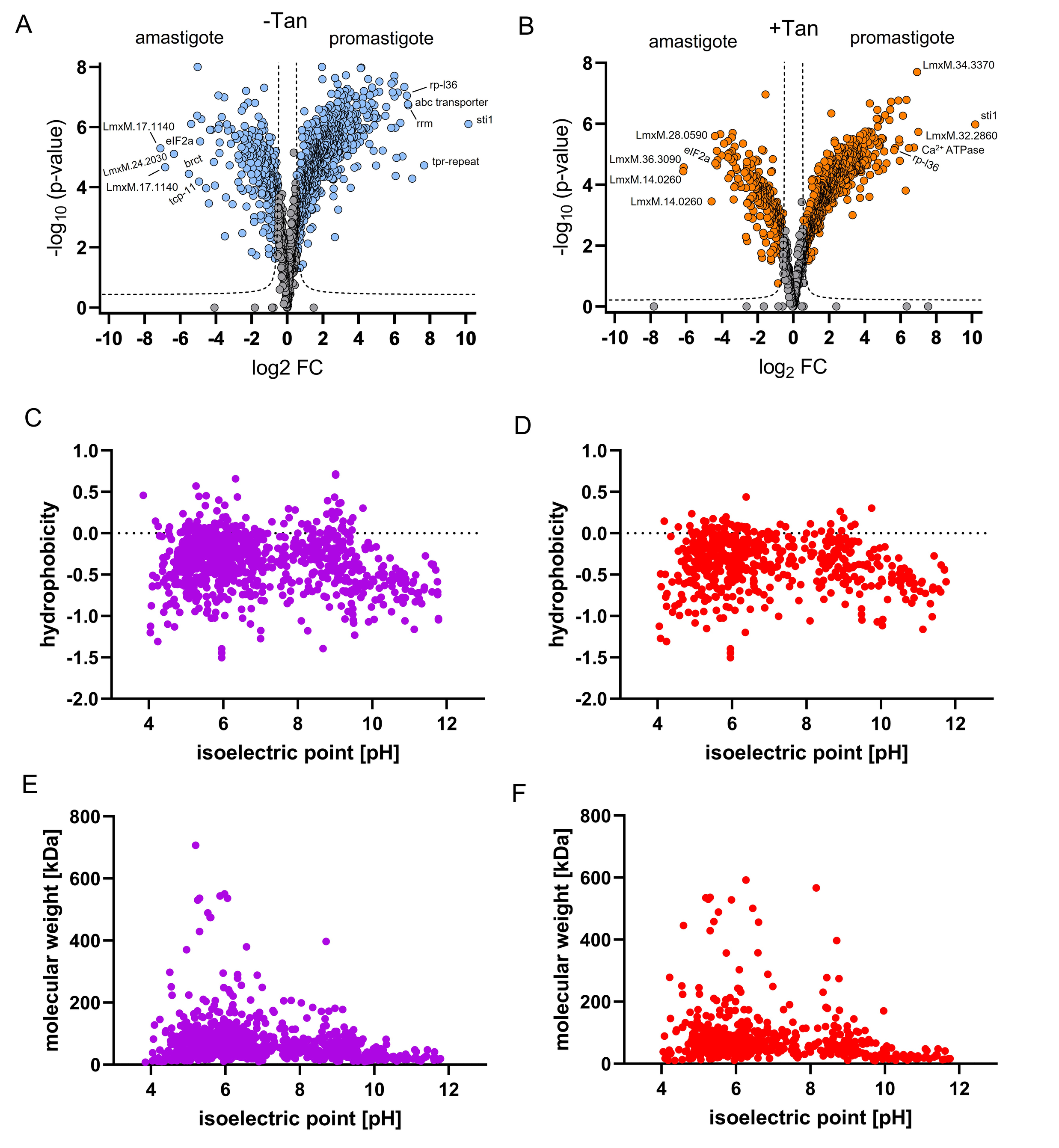
